# Phytochemical Screening and Antimicrobial Activity of *Ziziphus spina-christi* Stem Barks

**DOI:** 10.1101/2020.02.24.963157

**Authors:** Abdelrafie M. Makhawi, Mujahed I. Mustafa, Hajer A. Uagoub

## Abstract

This study was carried out in Khartoum State, during December 2017 The plant of Ziziphus *spina-christi* belong to family Rhamnaceae and locally known as Cedar, it was chosen for this study because of its using traditionally in treatment of many diseases. Microwave-assisted extraction (MAE) approach had been used to the dried sample at 170°C to reduce both time and extraction solvent volume, and to decrease the damage of bioactive compounds without extending the period of extraction. 20 g of sample was soaked with Petroleum ether, Ethyl-acetate, ethanol, methanol and distilled water for 72 hour. The extracts were concentrated using rotary evaporator at 40°C and were stored at 4°C. The phytochemical screening were carried out on different extracts of *Ziziphus spina-christi* stem bark and they showed to contain high amount of Tannins (4+ in all extracts), moderate amount of flavonoids and Triterpenes, trace amount of coumarins and Alkaloids and high amount of Saponins, Anthraquinones and Cardiac glycosides. The antimicrobial activity of extracts were evaluated against four standard bacteria species (gram positive; *Bacillus subtilis, Staphylococcus aureus*) and (gram negative; *Escherichia coli, Pseudomonas aeruginosa*; in addition of one standard fungi (*Candida albicans*). The results of antimicrobial tests indicated that the methanolic extract inhibited the growth of all microorganisms and most extracts showed several points of antimicrobial activity. These findings act as platform to assist in cure of bacterial and fungal infections.

## 1. Introduction

Medicinal Plants have been attaining great recognition all over the globe.[1, 2] They act as a very crucial therapeutic agents as well as valuable raw active compounds for manufacturing many traditional and modern treatments[3].

World Health Organization (WHO) recently reported that, there is an unprecedented increase in the occurrence of multidrug-resistant (MDR) infections worldwide [4, 5]. Currently, efforts are being focused to propose new drugs to multidrug resistant bacterial strains. Natural products, especially those obtained from medicinal plants, have demonstrated to be remarkable compounds with exceptional properties, such as rupturing the membrane of bacteria, enzymes activity inhibition and bacterial biofilm formation, which making them perfect applicants for these MDR bacteria[6-11].

*Ziziphus Spina-christi* is wide spread in tropical and subtropical region. The genus *Ziziphus Spina-christi* is widely distributed in the Middle East. Since ages extracts of *Ziziphus spina – christi* have been used as inflammatory treat toothache, analgesic, pectoral, astringent (LF), anti-rheumatic, purgative (FR), for stomach pain, anti-helminthic[12-14].

*Escherichia coli* is the main cause of infant mortality and diarrheal diseases. In the severe cases of diarrhea, the patient may have bloody diarrhea and can become life threatening. *E. coli* also responsible for urinary tract infection has become resistant to the drug that has been used to cure it [15, 16]. *Staphylococcus aureus* is resistant to antibiotics such as penicillin and methicillin; *Staphylococcus aureus* can cause life-threatening diseases such as sepsis and endocarditis; [17, 18] therefore, biochemical screening studies provide a promising alternative for drugs production from secondary metabolites of medicinal Plants due to its fewer side effects and cheaper cost.

This study focuses on determination of the chemical components and biochemical screening of *Ziziphus spina-christi* stems bark and to verify its antibacterial and antifungal activities.

## 2. Materials and Methods

### 2.1. Sample Collection

Fresh sample of *Ziziphus spina-christi* stem barks were collected from ALGAZIRA ISLANG, Northern of Omdurman, Khartoum State. The samples were washed and dried in open air in shade for two weeks.

### 2.2. Plants extract Protocols

Different extracts were prepared according to Robinson method [19] with little methodology development, as described: Microwave-assisted extraction (MAE) approach had been used to the dried sample at 170°C to reduce both time and extraction solvent volume, and to decrease the damage of bioactive compounds without increasing the period of extraction.[20] Twenty grams of sample was sopping with Petroleum ether, Ethyl-acetate, ethanol and methanol for 72 hours. The extracts were concentrated using rotary evaporator at 40°C, and finally the dried extracts were store at 4°C.

### 2.3. Preparation of aqueous extract

Fine powder (100gm) was poured with distilled water (300 ml) and left for 24 hours at room temperature. The mother liquor was filtered. The filtrate was vaporized till it reached dryness state; the residue thus obtained was the aqueous plant extract. All extracts were stored under aseptic conditions in bottles at 20 °C.

### 2.4. Phytochemical screening

#### 2.4.1. Test of alkaloids

A 3ml of extract was pour on petri dish and dried in water path, then dissolved in 10 ml of HCL 2% OR NH4OH 10% and transferred in three test-tube, each one contain 1ml, few drops of Dragendorff’s reagent were added into each tube (Dragendorff’s give orange precipitate. Wagner’s give reddish precipitate and Hager’s give yellow precipitate) which indicate the present of Alkaloids.

#### 2.4.2. Test of Flavonoids

A 2ml of extract evaporated on petri dish and then 10ml of ethanol were added, then transferred into four tests tube, the one added 1ml of NaOH that give yellow color, the second test tube Mg powder were added followed by 1drop of H_2_SO_4_ put in water path. The formation of a pink, crimson red which indicate the present of Flavonoids, the third test tube AlCl_3_ were added the formation of creamsh color indicated the present of Flavonoid, then ammonium solution were added the fourth test tubes. The formation of yellow/orange color indicated the present of Flavonoid.

#### 2.4.3. Test of Triterpenes and sterols

A 2ml of extract was dried in water path and dissolved in 6ml of chloroform, a few drops of concentrated sulfuric acid were added along the side of the test tube two layers was formed, the upper green color indicated the presence of sterol, and the lower red brown ring indicated the presence of triterpenes.

#### 2.4.4. Test of Tannins

A 2ml of extract poured in petri dish and late to dry, then was dissolved in 10ml ethanol solution and divided into two test tubes, the one 0.5ml of ferric chloride was added the formation of blue dark color indicator the presence of tannins, the second 0.5ml of gelatin salt (10%) was added, the formation of white precipitate point to the presence of tannins.

#### 2.4.5. Test of Saponins

0.5 ml of extract was poured in test tube, 10 ml of distilled water were added, the tube was strongly shaken for 30 seconds. The tube was allowed to stand and observed for creation of foam, which persisted for an hour, was taken as evidence for subsistence of saponins.

#### 2.4.6. Test of Coumarins

A 1g of plant powder was soaked in 10 ml distilled water in test tube and filter paper attached to the test tube to be saturated with the vapor after spot of 0.5N of potassium hydroxide put on it, then the filter paper was examined under Ultra Violet (UV) light, the presence of coumarins was designated if the spot have found to be adsorbed the UV light.

#### 2.4.7. Test of Cardiac glycosides

A 0.1g of plant powder was soaked in 1ml glacial acetic acid with one drop of ferric chloride solution, 1ml the sulphuric acid was added under layer, a brown ring obtained was indicated the presence of glycosides.

#### 2.4.8. Test of Anthraquinones

A 0.1g of plant powder was dissolved in 1ml water and 5ml of chloroform was added and shacked for 5 minutes, after shaking two layer were formed, the chloroform layer was separated. 1ml of ammonia solution (10%) was added into 1ml of chloroform the appearance of pink or red or violet color designates the presence of Anthraquinones.

### 2.5. Microbial analysis

#### 2.5.1. Preparation of Standard Microorganism’s Suspensions

Preserved bacterial strains (obtained from the Department of Microbiology, University of Bahri, Khartoum, Sudan) were cultured on nutrient broth. Full loop of bacteria was inoculated in nutrient broth at 37°C in incubator for 24 hours; the growth was observed in the broth, and then sub-cultured to nutrient agar plates for 24 hours. When growth confirmed as the correct species by examining it under a light microscope (DM3000; Germany), the individual colonies were again sub-cultured to nutrient agar slopes and inoculated at 37°C in incubator until the growth occurred. 1 ml of the bacteria suspension was diluted with 9 ml of sterile saline that was taken with pipette and shaken gently to produce a suspension that containing about (10 −10 CFU/ml). The suspension was stored in the refrigerator at 4°C till used. Preserved fungal strains were cultured on Saboraud dextrose agar, prior to antifungal susceptibility screening [21].

#### 2.5.2. Extracts analysis for antimicrobial activity

The method used to test the sensitivity to the plant extract, as outlined by the International Commission on Microbiological Specifications for Foods [22]. 0.1 ml of a broth culture of each bacterium (*Escherichia coli, Staphylococcus aureus, Bacillus subtilus and_Pseudomonas aeruginosa*) and fungi (*Candida albicans*) was inoculated on each plate. Wells were done under aseptic conditions. Each well was filled with 20 μl of stem bark extract of *Ziziphus spina –christi* of different solvents at different concentrations (100, 50, 25 µg/ml), then plates were cooled for 2 hours to allow proper diffusion before incubation at 37°C for 48 hours. After incubation, the inoculated sensitivity plates were removed from the incubator and inhibition zones around the wells were measured. An inhibition zone measuring more than 14 mm was considered sensitive.[23]

## 3. Results

### 3.1. Results of extraction and physical properties

Five solvents were used in successive polarities to extract secondary metabolites from *Ziziphus spina –christi* stem barks and their properties.

The results of the extractive values of *Ziziphus spina-christi* as following: for methanol 4.43% (crystal bright black), followed by ethanol 4.4% (crystal bright black), ethyl acetate 2.885% (brown powder), petroleum ether 1.9% (brown crust fine powder), and distilled water 3.85% (brown powder). The extraction results and fractionation yields are shown in (Table 1).

**Table (1):** National Collection of Type Culture (NCTC), Colindale, England:

| Name of standard organism | ATCC code | Type of organism |
| --- | --- | --- |
| *Escherichia coli* (E. c) | 25922 | Gram negative bacteria |
| *Pseudomonas aeruginosa* (P. a) | 27853 | Gram negative bacteria |
| *Bacillus subtilis* (B. s) | 8236 | Gram positive bacteria |
| *Staphylococcus aureus* (S. a) | 25923 | Gram positive bacteria |
| *Candida albicans* | 7596 | Fungi |

(ATCC): American Type Culture Collection Rockville, Maryland, USA.

### 3.2. Phytochemical screening activity of *Ziziphus spina –christi* stems barks

Phytochemicals analysis carried out on the *Ziziphus spina –christi* stems barks extracts unmasked the presence of bioactive compounds, which qualitatively analyzed and presented in **(Table 2)**.

**Table (2):** Properties and extractive value of *Ziziphus spina –christi* stem barks:

| Solvents of extraction | Characteristic | Color | Weight (g) | Yields (%) |
| --- | --- | --- | --- | --- |
| Methanol | crystal | Bright black | 8.86 | 4.43 |
| Ethanol | crystal | Bright black | 8.8 | 4.4 |
| Ethyl acetate | powder | brown | 5.77 | 2.885 |
| Petroleum ether | Crust of fine powder | brown | 3.8 | 1.9 |
| Distilled water | powder | brown | 7.7 | 3.85 |

### 3.3. Antimicrobial tests

The extracts of *Ziziphus spina –christi* stem bark at concentrations (100mg/ml, 50mg/ml, 25mg/ml), were subjected to antimicrobial tests by using cup plate agar diffusion method and inhibition zone were measured in (mm) against four bacterial strains and one fungi.(**Table 1**) The range of inhibition was found 14-25 mm.

## 4. Discussion

Sudan is the largest country in Africa with rich plants due to the climate diversity which are used in traditional medicine to treat several diseases.[24-26] In the present study, five solvents were used in successive extractable method from Ziziphus *spina-christi* stem barks, methanol solvent gave generally higher extractability than those obtain from (ethanol, ethyl acetate, distilled water and petroleum ether) (**Table 2**). In case of ethyl acetate, distilled water the consistency of extractable material seem to be brown powder, with differences in yield, and the extract of ethanol and methanol seem to be crystal bright black with differences in yield. Petroleum ether it is lower extractability than all other it is crust of fine brown powder. Variation was observed in the colors of extracts may be were reflection to type of solvent ingredients’ of plant.

Phytochemical screening of chemical constituents of *Ziziphus spina-christi* stem bark extracts, were found that all with high amount of tannins. The flavonoids were present in moderate amount in methanol and ethanol extracts. Moreover, all extracts found contain low amount of coumarins; while alkaloids present in low amount in ethanol. triturbin and sterol present in moderate amount in all extracts. In addition cardic glycoside was found in a high amount in all extracts. Extracts showed to contain high amount of Saponins. The extracts showed to contain high amount of anthraquinones;(**Table 3**) all act as antioxidants, as well known, antioxidants decrease free radical attack on DNA and therefore prevent mutations that cause cancer;[27] this difference may due to the condition of the experiments or well as in differences in methodology, or/and in variation of amount of secondary metabolites. In contrast synthetic antioxidants have recently been shown to cause critical health issues such as liver damage, due to their carcinogenicity effect. Hence, the development of safer antioxidants from natural sources has been recognized, and medicinal plants have been used as a reliable source of alternative medicines to treat various diseases.[28]

**Table (3):** Shows phytochemical screening results of *Ziziphus spina-christi* stem barks:

| Test/ Solvent | Methanol | Petroleum ether | Distilled water | Ethyl acetate | Ethanol |
| --- | --- | --- | --- | --- | --- |
| NaOH | +++ | ++ | ++ | ++ | +++ |
| AlCl<sub>3</sub> | +++ | + | + | ++ | +++ |
| NH<sub>4</sub>OH | +++ | + | ++ | ++ | ++ |
| Mg/H<sub>2</sub>SO<sub>4</sub> | ++++ | + | +++ | ++ | ++++ |
| Tannin Ferric chloride | ++++ | ++++ | ++++ | ++++ | ++++ |
| Gelatin | ++++ | ++++ | ++++ | ++++ | ++++ |
| Wagner's | + | + | + | + | ++ |
| Dragendorff's | + | + | ++ | + | ++ |
| Hager's | +++ | + | ++ | +++ | +++ |
| Lieberman | ++ | ++ | + | ++ | ++ |
| Salkowski | +++ | +++ | ++++ | +++ | +++ |
| Coumarins KOH/UV | + | — | + | ++ | + |
| Saponins Foam test | ++++ | + | ++ | + | ++++ |
| Anthraquinones | ++++ | + | + | ++++ | ++++ |
| Cardiac glycoside | ++++ | ++++ | +++ | ++++ | +++ |

(++++): high amount, (+++): moderate amount, (++): low amount, (+): trace amount, (-): not found.

The methanol showed high activity at all concentrations (100%,50%,25%), against the clinical isolation of E. coli (22,21,20) respectively, as well as against of *Pseudomonas arginosa* (24,23,19), *Staphylococcus aureus* (21,20,18), and showed low activity against *Candida albicans* (20,18.17), and showed high activity against *Bacillus subtitles* (25,22,20) (**Figure 1**). Ethanol was showed against these bacteria: E. coli (20,18,17), *pseudomonas arginosa* (19,17,16), *Staphylococcus aureus (21,20,18), and showed as high amount against Bacillus subtitles (22,20,18),and at low amount against Candida albicans (20,18,15). The ethyl acetate extracts high activity against Bacillus subtitles (22, 20, 18), and low activity against Candida albicans (17, 16, 15), E. coli (18,17,15), Pseudomonas arginosa (15, 14, ND), and have no any activity against Staphylococcus aureus. Distilled water showed high activity against Bacillus subtitles (24,23,20), and low activity against Candida albicans (18,17,16), Staphylococcus aureus(18,16,14), Pseudomonas arginosa (20,17,15), Staphylococcus aureus*(16,15,14), (**Figure 4**) while petroleum-ether has no inhibition zone against all tested microorganisms (**Table 4**).

**Table (4):** results of antimicrobial activities of *Ziziphus spina-christi* stem barks:

| Extract | Concentration | C.a | B.s | S.a | P.a | E.c |
| --- | --- | --- | --- | --- | --- | --- |
| Methanol | 100 | 20 | 25 | 21 | 24 | 22 |
|  | 50 | 18 | 22 | 20 | 23 | 21 |
|  | 25 | 17 | 20 | 18 | 19 | 20 |
| Ethanol | 100 | 20 | 22 | 21 | 19 | 20 |
|  | 50 | 18 | 20 | 20 | 17 | 18 |
|  | 25 | 15 | 18 | 18 | 16 | 17 |
| Ethyl acetate | 100 | 17 | 22 | – | 15 | 18 |
|  | 50 | 16 | 20 | – | 14 | 17 |
|  | 25 | 15 | 18 | – | – | 15 |
| Distilled water | 100 | 18 | 24 | 18 | 20 | 20 |
|  | 50 | 17 | 23 | 16 | 17 | 18 |
|  | 25 | 16 | 20 | 14 | 15 | 17 |
| Petroleum ether | 100 |  |  |  |  |  |
|  | 50 |  |  |  |  |  |
|  | 25 |  |  |  |  |  |

*B.s:* *Bacillus subtilis*; *S.a:* *Staphylococcus aureus*; *E.c,* *Escherichia coli*; *P.a:* *Pseudomonas aeruginosa*; *C.a:* *Candida albicans*; concentration of extracts 0.2ml/ cup (100 ,50,25mg/ml).

**Figure (1):**
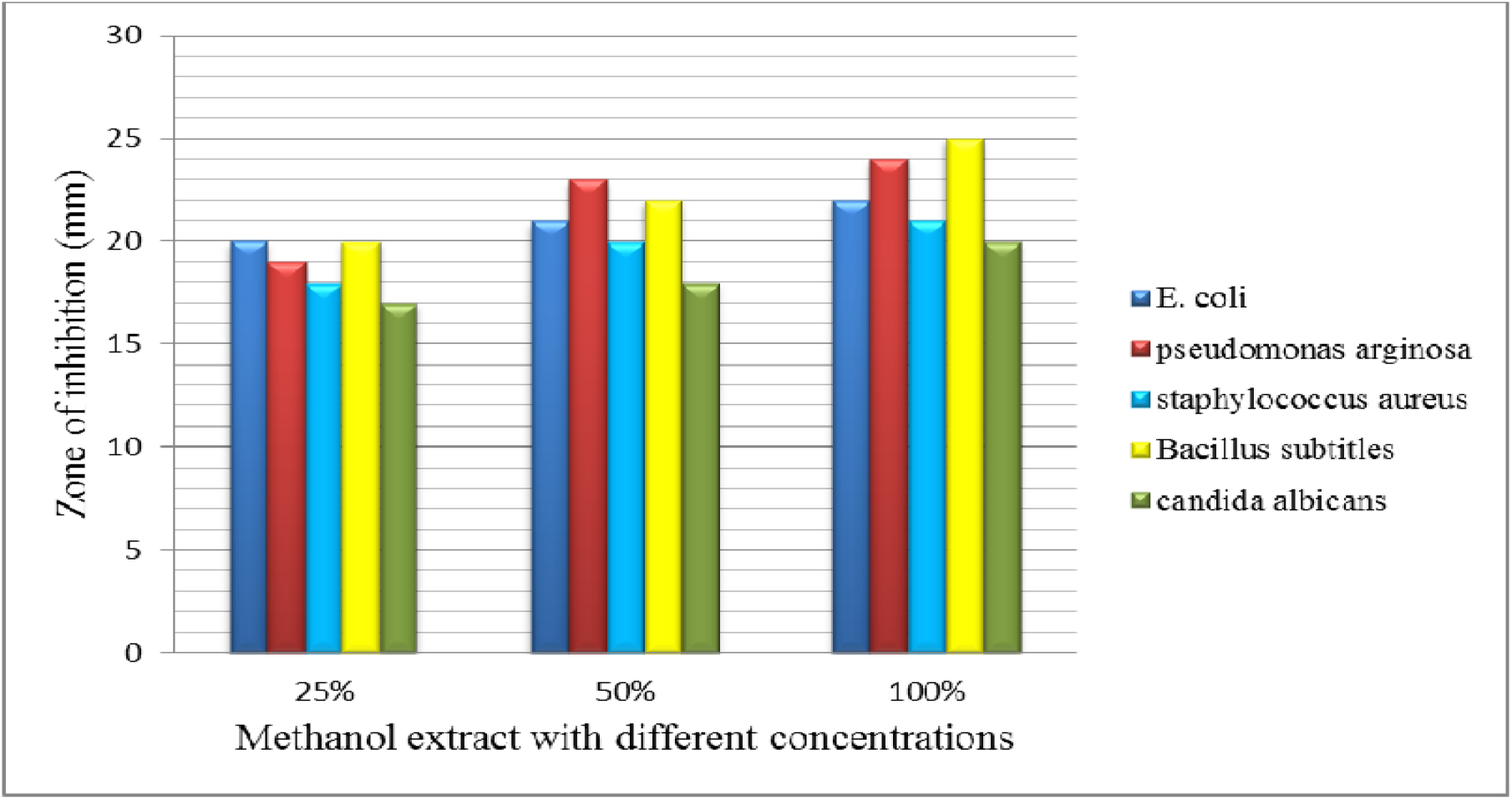
Inhibition Zone (mm) of bacterial and fungal pathogens by stem barks extract of *Ziziphus spina –christi*.

**Figure (2):**
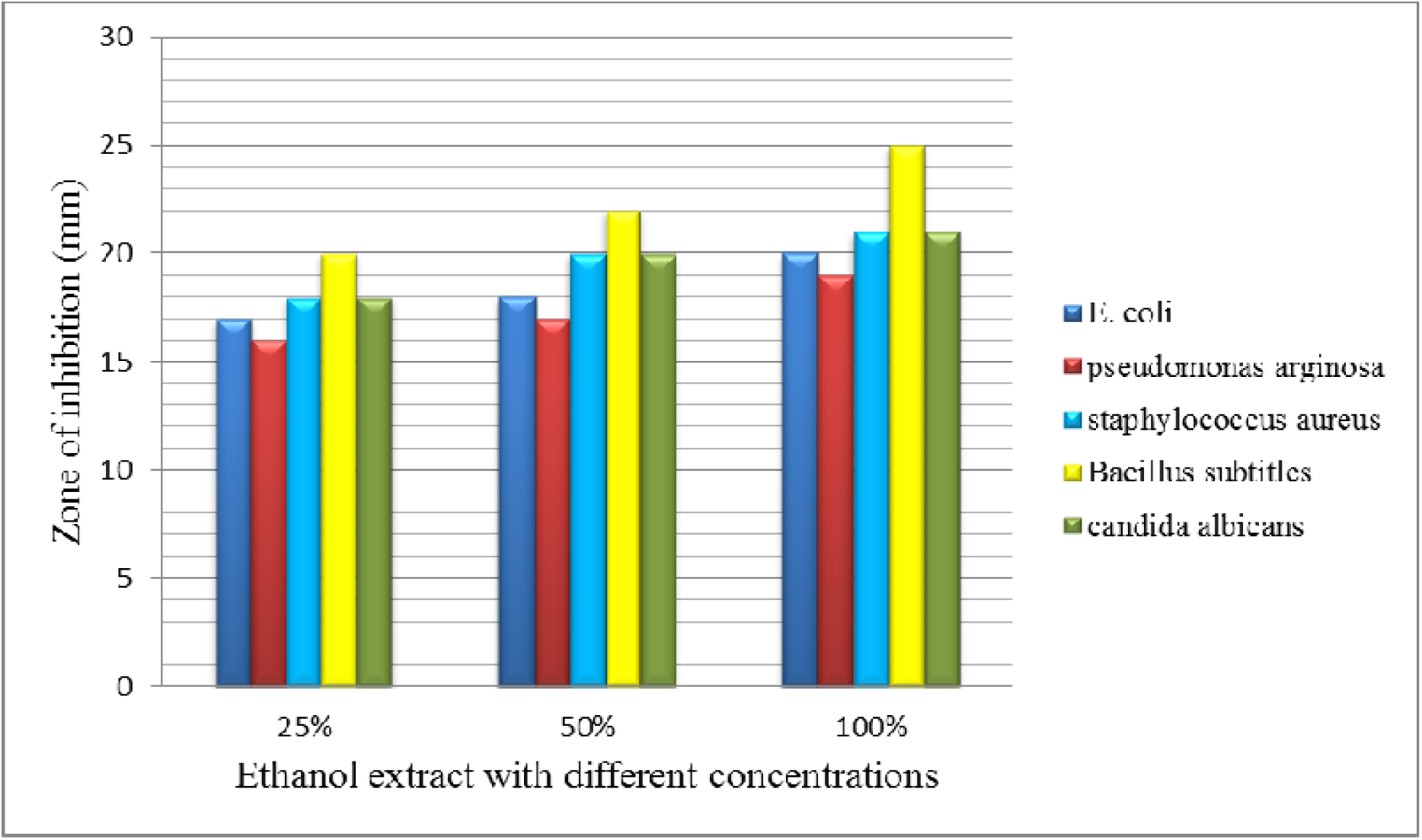
Inhibition Zone (mm) of bacterial and fungal pathogens by stem barks extract of *Ziziphus spina –christi*.

**Figure (3):**
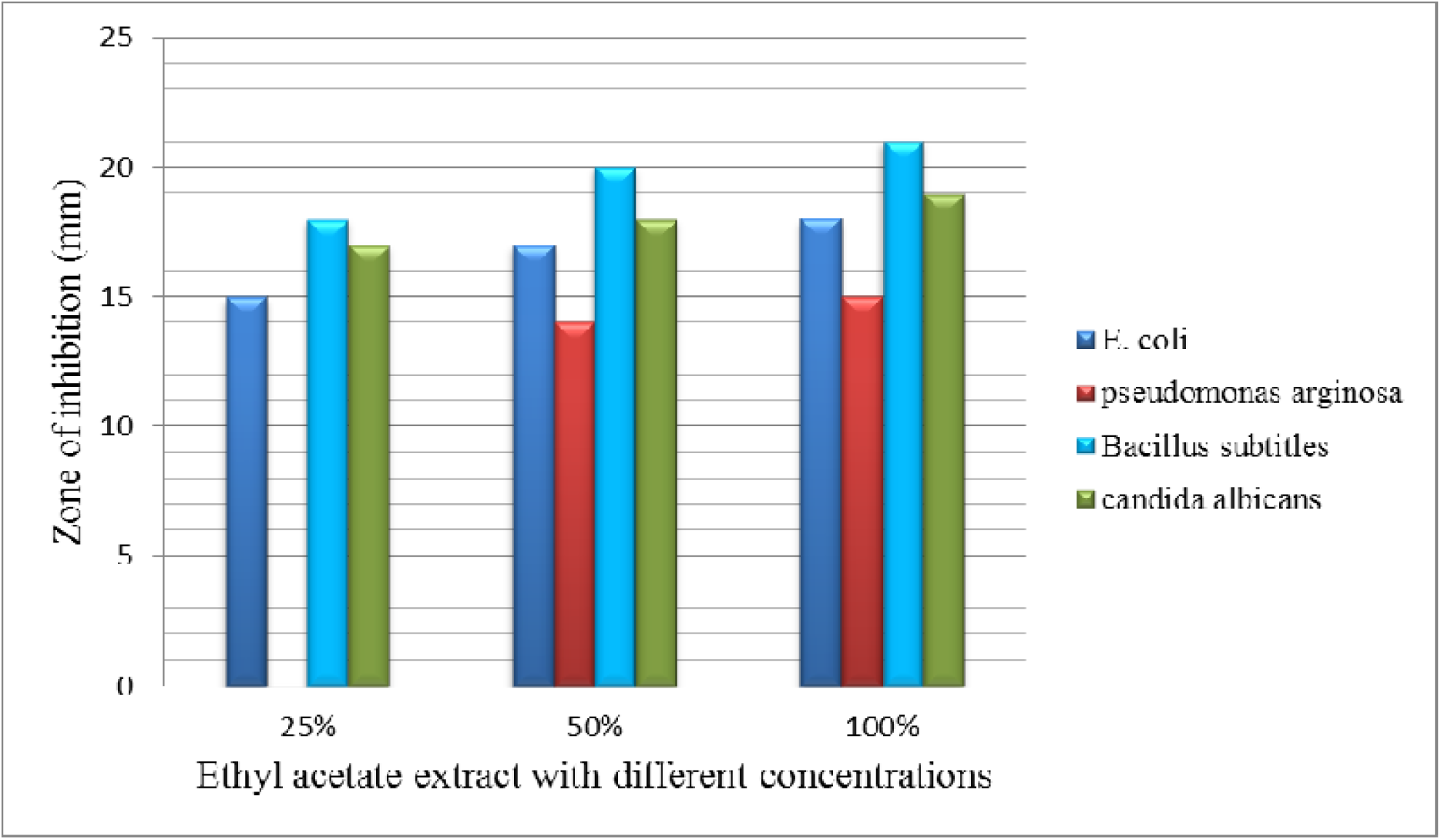
Inhibition Zone (mm) of bacterial and fungal pathogens by stem barks extract of *Ziziphus spina –christi*.

**Figure (4):**
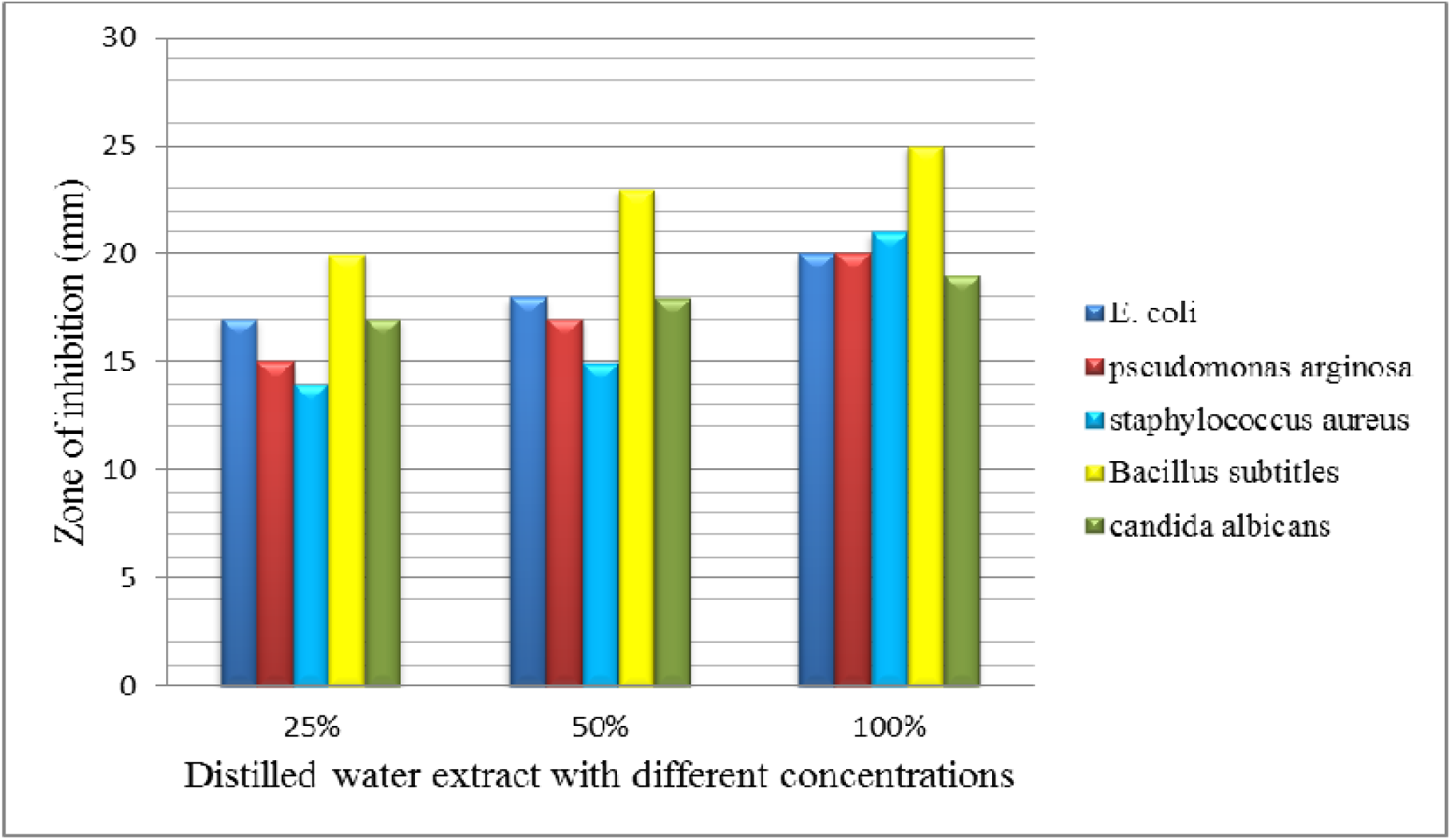
Inhibition Zone (mm) of bacterial and fungal pathogens by stem barks extract of *Ziziphus spina –christi*.

This compared with assays reported by El-kamali and Mahjoub[29] methanol extract of *Ziziphus spina–christi* stem bark was found effective against all tested Gram negative and Gram positive bacteria inhibition zone (range between 20-30 mm). Ethanol extract of *Ziziphus spina–christi* stem barks was found effective against all tested bacteria except Escherichia coli. Where Ethyl acetate of *Ziziphus spina–christi* stem barks was active against both Gram positive and Gram negative bacteria. Aqueous extract of *Ziziphus spina–christi* stem barks was moderately active against Gram positive and Gram negative bacteria inhibition zone (range between 15-17 mm). Petroleum ether extract of *Ziziphus spina–christi* stem bark had the ability to inhibit *Bacillus subtilis* (15 mm) and *Escherichia coli* (20 mm).

Our findings can be used to enhance the potential of currently used antimicrobial agents.[30] Further studies needed to ensure the safety of Ziziphus *spina –christi* stem bark and development of safer antioxidants from this natural source.

## 5. Conclusion

The presented results offer supporting evidence for effective use of selected plant extracts. Antimicrobial resistance is reported to be on the increase due to gene Mutation of the disease pathogens. *Ziziphus spina* □*Christi* was chosen For this study because of their reputation in folklore medicine as antimicrobial agents and usage of different parts in many diseases. Phytochemical screening was carried out and lead to presence of some Secondary metabolites the plant was showed to contain alkaloids, flavonoids, tannins, triterpenes, coumarins, saponins, cardic glycoside, anthraquinone, all act as antioxidants. The crude extracts were subjected to antimicrobial assays using cup Plate diffusion method and the inhibition zone were measured in mm. The methanol extracts of *Ziziphus spina-christi* gave good results activities against bacterial and fungal organisms were used, while the petroleum showed absence of inhibition zone against four bacterial and on fungal used.

## Data Availability

The data which support our findings in this study are available from the corresponding author upon reasonable request.

